# Low-resolution FAIMS for increased peptide coverage in low-load and single-cell proteomics

**DOI:** 10.1101/2025.08.22.671812

**Authors:** Dominic G. Hoch, Manuel Matzinger, Karl Mechtler

**Author notes:** These authors contributed equally (Dominic G. Hoch, Manuel Matzinger).

## Abstract

Field Asymmetric Ion Mobility Spectrometry (FAIMS) enhances signal to noise ratio filtering ions based on their differential mobility, making it an indispensable tool for single-cell and low-input proteomics. Here, we investigate the impact of FAIMS resolution tuning via electrode temperature modulation to improve peptide identification sensitivity. We demonstrate that lowering FAIMS resolution broadens the compensation voltage window, thereby increasing ion transmission. This “low-resolution” mode significantly improves peptide identifications from a concentration range of HeLa digests by up to 18%. A weaker but still beneficial effect in peptide identifications and enhanced sensitivity could also be reproduced for single-cell samples. This leads to proteomic fingerprint shifts, resulting in distinct populations in principal component analysis from the very same cell-type in dependence on FAIMS resolution. Moreover, the increased ion counts of runs employing “low-resolution” FAIMS improve the quantitative precision of low-load measurements. These findings offer a practical optimization strategy for FAIMS-based low-input proteomics workflows, that allow for improved results by changing a single setting within the MS method without the need of any change in hardware adoption or data analysis pipelines.

## Introduction

Field Asymmetric Ion Mobility Spectrometry (FAIMS) separates gas-phase ions based on their mobility differences under alternating high and low electric fields at atmospheric pressure. Ions are transported by carrier gas flow between two closely spaced electrodes, with an asymmetric electric field waveform applied orthogonally to this ion flow.^1,2^ This waveform alternates between a high-voltage, short-duration pulse of one polarity and a lower-voltage, longer-duration pulse of the opposite polarity. The high-field segment is referred to as the dispersion voltage (DV).

Ion mobility varies with electric field strength, causing ions to behave differently during the high and low field phases. Ions with equal mobility in both fields experience no net displacement and pass through the device. Conversely, ions with differing mobilities drift toward one electrode, potentially colliding with it and being neutralized and removed from the ion stream. To selectively transmit ions of interest, a small direct current compensation voltage (CV) is applied to counterbalance the net drift. By adjusting the CV, only ions with specific mobility characteristics are allowed to pass through to the detector, while others are filtered out. This continuous filtering process allows FAIMS to separate ions based on their differential mobility, enhancing selectivity and reducing chemical noise prior to mass spectrometric analysis. FAIMS is often employed as an additional filtering step between liquid chromatography and mass spectrometry, improving the selectivity and sensitivity of various applications like bottom-up proteomics, ^3,4^ targeted proteomics,^5^ top-down proteomics,^6–8^ PTM analysis,^9^ metabolomics,^10,11^ etc.

FAIMS has been extensively utilized in proteomics to improve the identification and quantification of proteins and peptides. By reducing chemical noise and enhancing selectivity, FAIMS contributes to more accurate and sensitive proteomic analyses. This is particularly beneficial in complex biological samples where the dynamic range of protein concentrations can span several orders of magnitude. In single-cell proteomics, where the amount of available material is extremely limited, FAIMS plays a crucial role in enhancing analytical performance.^12–15^ FAIMS reduces background noise and chemical interferences, allowing for the detection of low-abundance proteins that might otherwise be missed.

Despite all these advantages, there are limitations to the usage of FAIMS in mass-spectrometry based analyses. The post-switching delay time that is required with every change of compensation voltage limits the number of chosen CVs per experiment. In recent years, in the context of single-cell proteomics, only one CV is used during fast gradients that are required to have reasonable throughput. This significantly increases the signal-to-noise ratio of the analytes that pass the electrode assembly and enter the mass spectrometer, but it filters out other peptides that have a different collisional cross section (CCS) and are not transmitted to the MS with a chosen CV. Choosing more CVs per experiment would increase the peptide coverage per experiment, but the acquired datapoints per peak and per CV-fraction will be reduced by the number of CVs, negatively impacting the quantitative precision of measurements.

The temperature of the electrodes significantly influences ion mobility in several ways: Elevated electrode temperatures can promote thermal desorption of ions that are weakly bound to surfaces or contaminants, thereby improving the purity and clarity of the ion stream. Maintaining stable electrode temperatures ensures consistent ion mobility measurements, leading to reproducible results across different runs and samples.^1^ Higher temperatures generally increase the kinetic energy of the ions, which can lead to changes in their mobility. Temperature control can be used to optimize the separation of ions with similar mobilities, improving the resolution of the FAIMS device. The temperature of the inner electrode can be lowered to increase the resolution in the CV dimension. This approach has been used in targeted analysis to increase selectivity of isobaric molecules. To increase the fraction of ions per CV that pass through the electrode assembly to the MS, we sought to decrease the FAIMS resolution by lowering the outer electrode temperature.

## Results

### Evaluation of FAIMS resolution effect using selected ions in a static spray

Firstly, we evaluated if the resolution in CV dimension changes upon changing the electrode temperatures. We directly injected Pierce™ FlexMix™ Calibration Solution (Thermo Fisher Scientific) at 5 µL/min into the mass spectrometer using a syringe pump with FAIMS attached and determined the optimum CV values for the calibration mix ions hexamethoxphosphazene (322.0481 m/z), MRFA (m/z 524), Hexakis(2,2-difluoroethoxy)phosphazene (622.0289 m/z) and Hexakis(2,2,3,3-tetrafluoropropoxy)phosphazene (922.0098 m/z) for different outer electrode temperatures (Figure 1 A-D). Using FAIMS in standard resolution mode (in: 100°C, out: 100°C) showed narrower peaks compared to choosing a “low-resolution” setting (in:100°C, out: 80°C). With the “low-resolution” mode, the ions pass the FAIMS and enter the MS over a wider CV range (Figure 1 E). The change in resolution further results in a slight shift in the apex CV towards more positive values (**Figure 1** F). Of note, for some ions, namely 524 m/z and 622 m/z, (**Figure 1** B & C) the ion transmission seemed further improved by lowering FAIMS resolution as indicated by higher a higher TIC observed. This is of potential advantage for low input or single cell proteomic applications as well.

**Figure 1:**
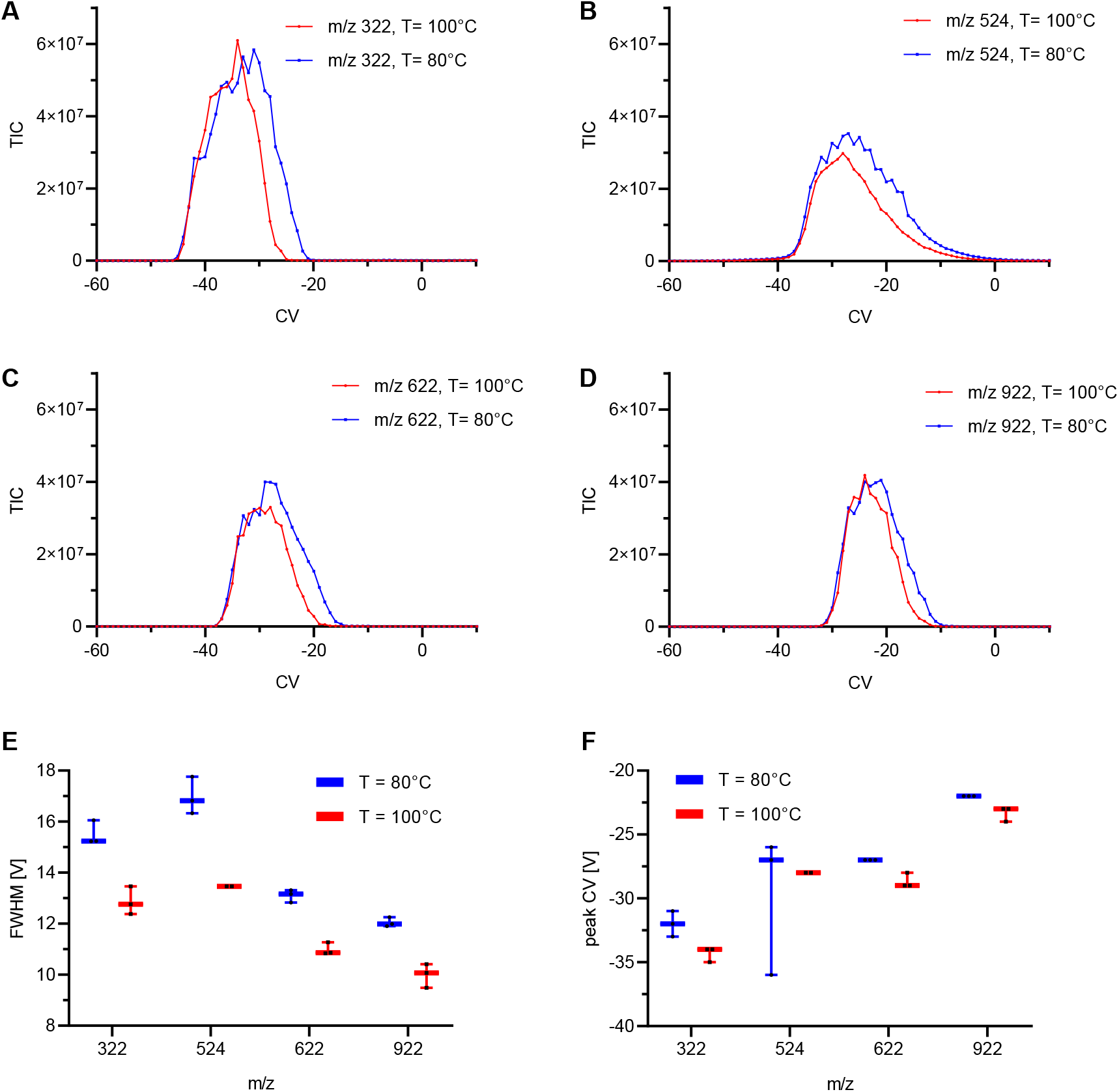
FAIMS resolution assessed by FlexMix injection. Total ion chromatograms from SIM scans upon FlexMix injection with standard resolution (T_outer_ electrode = 100°C) or low resolution (T_outer_ electrode = 80°C) when performing a CV scan for m/z 322.0481 (**A**), 524.0000 (**B**), 622.0289 (**C**) or 922.0098 (**D**), respectively. A representative replicate out of 3 technical replicates is shown. **E**: Box plots show the full width half maximum (FWHM) of peaks resulting from scanning thorough CVs. **F**: The peak apex CV is shown. All box plots within this figure show median values with error bars depicting standard deviations, n=3.

### Impact on identification numbers and data quality

To determine the effect of the FAIMS electrode temperatures on the data quality, we injected 250pg HeLa digest using a 50SPD gradient and varied the temperatures of the outer electrode temperature from standard 100°C to 80°C in 5°C steps. The protein identification did not change significantly and did not show a trend when changing the electrode temperatures (

**Figure 2A**). Interestingly, the peptide identifications gradually increased with decreasing outer electrode temperatures. The largest effect was observed using an outer electrode temperature of 80°C: An average of 28,063 peptides in standard vs. 35,258 peptides with the lowest electrode temperature (

**Figure 2:**
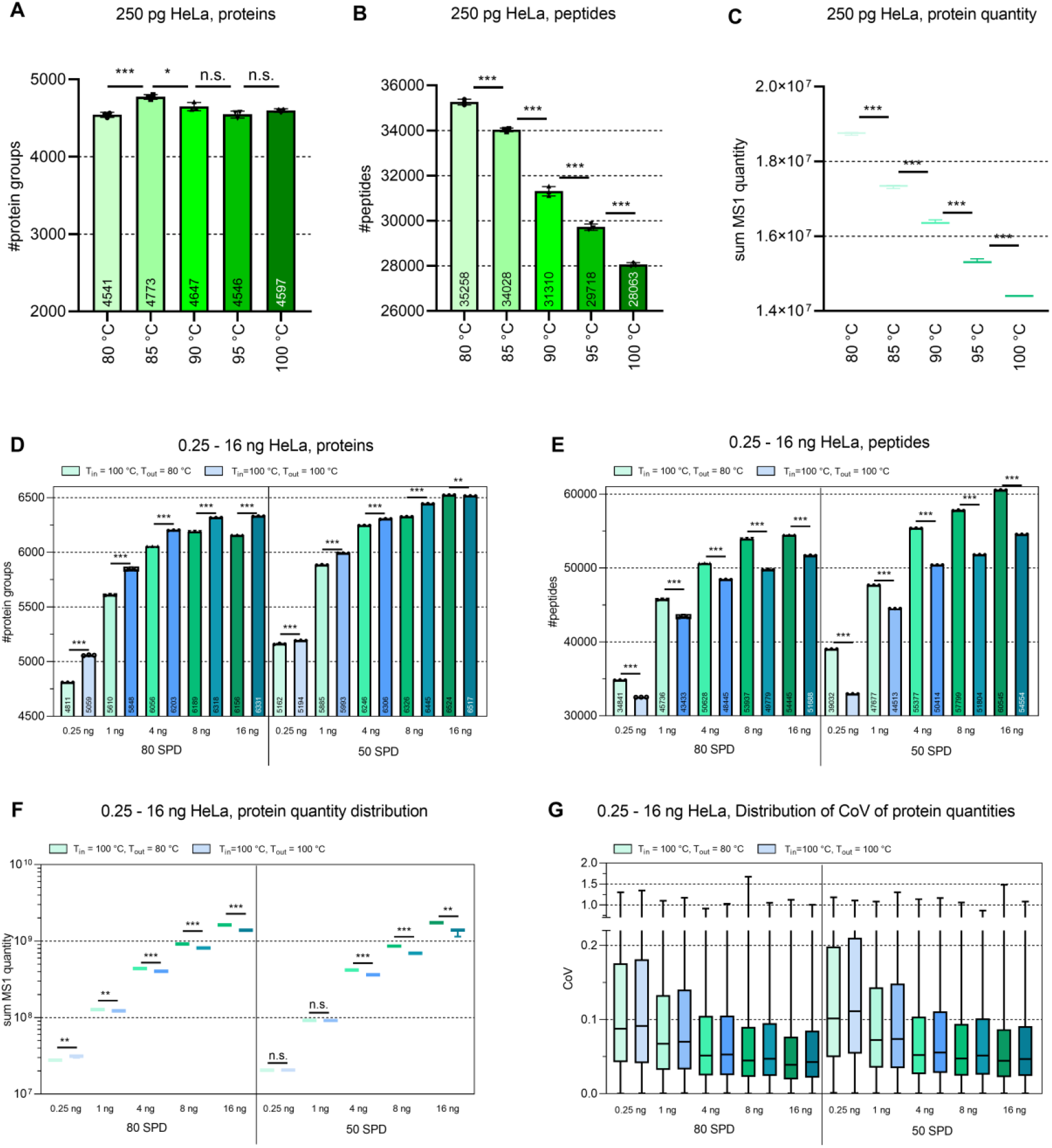
Influence of FAIMS electrode temperature on protein and peptide identifications and their quantities. Data shown from diluted HeLa digests injected in quantities as indicated. Bars show average values, error bars indicate their standard deviations, n = 3 technical replicates. For statistical comparison of two average values unpaired two tailed students t tests were performed, with *** for p<0.001, ** for p<0.01, * for p <0.05 and n.s. for p ≥ 0.05. **A**: Proteins identified within 250pg HeLa, **B**: Peptides identified from 250pg HeLa, **C**: Average summed of protein quantity, with quantification performed on MS1 level, with data in **A-C** using a 50 SPD gradient. **D**: Proteins identified within 0.25 – 16 ng HeLa digest using a 50 or 80 SPD gradient as indicated, **E**: Peptides identified within 0.25 – 16 ng HeLa digest, **F**: Box plots showing the distribution of total protein quantity and **G**: Box plot showing the distribution of coefficient of variations from protein quantities from proteins quantified in n=3 replicates. Error bars indicate the minimal to maximal value while lines within boxes indicate median values.

**Figure 3:**
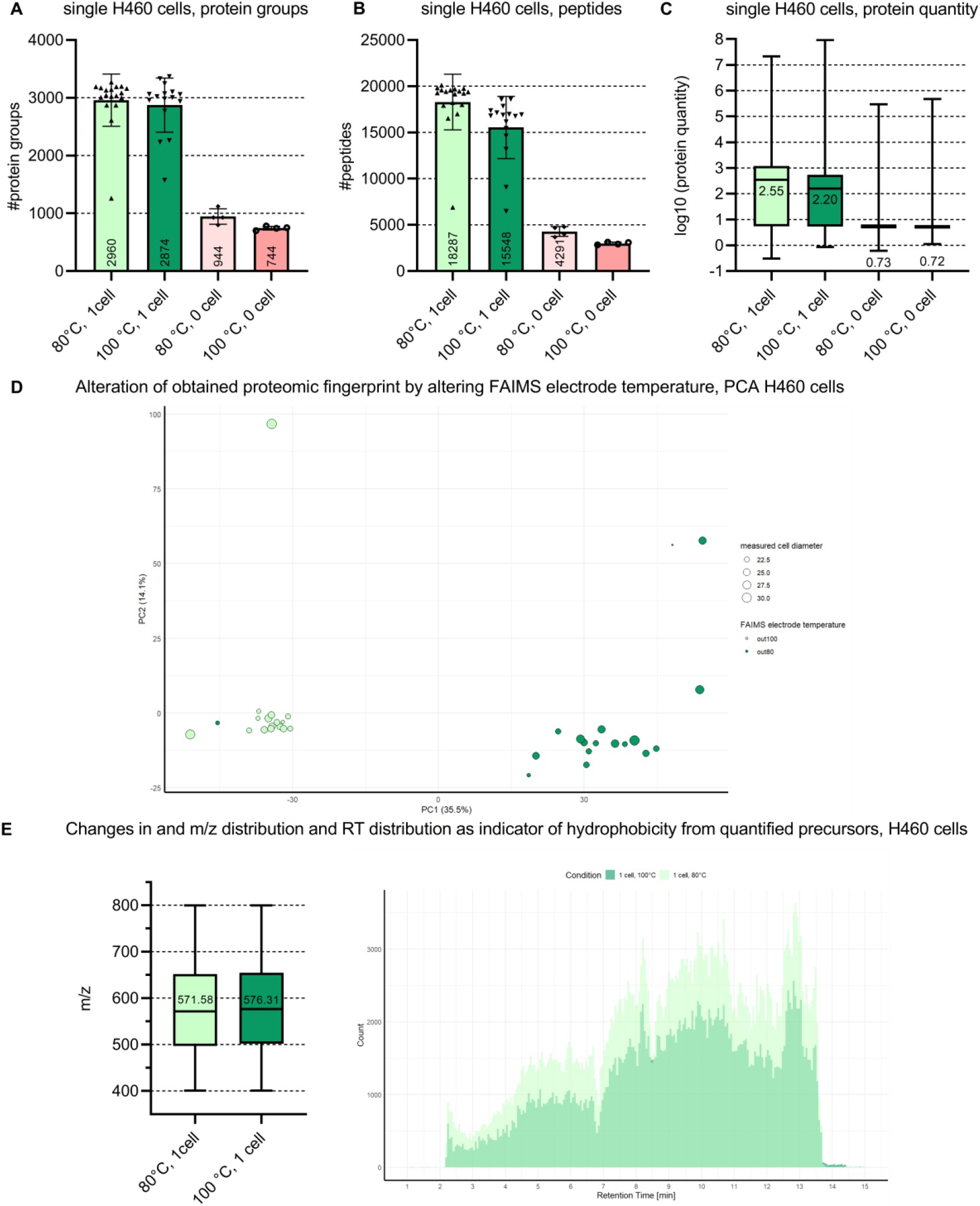
Analysis of single H460 cells using high or low FAIMS resolution. Bars show the average number of quantified protein groups (**A**) or peptides (**B**) from single cells respectively; error bars indicate standard deviations and values from individual cells are shown as black dots. Box plots in (**C**) show the distribution of protein quantities in single cells and 0 cell negative controls respectively. Missing values were filtered; data was not normalized. N=16 cells for 10°C, 18 cells for 80°C and n=4 for 0 cell controls. **D**: PCA with each dot representing a single H460 cell measured at a given FAIMS electrode temperature. Dot sizes represent measured cell diameter as indicated. **E**: Left: Boxplot showing the distribution of m/z values of all quantified peptide groups and right: Histogram of quantified precursors in all H460 cells measured at the given FAIMS electrode temperature over the entire retention time and using 250 bins.

**Figure 2B**). Moreover, we observed an increased summed protein quantity, while gradually decreasing the outer electrode temperature (

**Figure 2C**). We also determined a higher total LC peak fragment ion count when using “low-resolution” FAIMS in 250pg HeLa 50SPD runs compared to the standard settings (**Supplemental** Figure 1).

We observe a clear beneficial trend with lowered outer electrode temperatures that seems to potentially continue for temperatures lower than 80°C. However, when defining 75°C or less as outer electrode temperature we could not reach and hold that temperature stably. This is likely reasoned by a too big temperature difference between inner and outer electrode with the cooling gas flushing the 100°C hot inner electrode first not being able to maintain 75°C or less for the outer electrode. We therefore kept the minimal outer temperature at 80°C which we were able to reproducibly maintain stable over the entire gradient (**Supplemental Figure**).

Encouraged by these results, we investigated if there is a dependency on the loaded peptide amount. We therefore ran a concentration range of HeLa digest (250 pg, 1 ng, 4 ng, 8 ng, 16 ng) to compare the standard electrode temperature setting and the lowest outer electrode temperature using two gradients, 50SPD and 80SPD. Here again, we observed a significant effect on the peptide identifications for all tested loads and both gradients, when lowering the outer electrode temperature to 80°C (

**Figure 2E**). The increase in peptide identifications ranged from 6-18% for the 50SPD gradient and 4.8% - 8% for the 80 SPD gradient (see supplemental Error! Reference source not found.). On the protein level, the same effect as previously was observed: the protein groups are slightly lower for both gradients but never decreased more than 5% (

**Figure 2D**). The biggest negative effect was observed for the 250pg load at 80SPD – 5059 IDs vs. 4811 IDs. The summed MS1 quantity was also consistently higher when decreasing the outer electrode temperature with the one exception of the 80SPD 250pg HeLa run (

**Figure 2F**). In line with these higher quantities and the higher median contribution to the total LC peak fragment ion count (Supplemental Figure 1), our data also suggests a better quantitative precision of measurements using “low-resolution” FAIMS. This is indicated by slightly lowered Coefficient of Variation (CoV) values over the entire range of tested input amounts and gradient lengths (**Figure 2G**).

### Impact of FAIMS resolution on Single Cell Proteomic Samples

We moved forward by analyzing single cells to determine if the electrode temperature also positively influences the measurement of actual single cells. H460 cells were prepared using our previously published OnePot workflow^16^ and measured using the 80SPD gradient. In the case of single cells, there was no significant change in identifications observed, neither on the protein nor the peptide level, with a positive trend towards 80°C on peptide level again though. This is also true for the distribution of protein quantities again at higher levels for the “low-resolution” mode (Error! Reference source not found. **A-C**). Of note, we included 0-cell controls in our analysis that contain identical components to single cell samples and were processed in the exact same way on the same 384-well plate but with no cell isolated into the respective well. This indicates the level of background signals obtained from contaminations and the added protease itself. Lowering the resolution hence slightly improves sensitivity, crucial for single cell proteomic applications. This yields differences in the obtained proteomic fingerprint leading to distinct populations of cells within a PCA plot depending on the electrode temperature used (Error! Reference source not found. **D)**. While improved sensitivity is the most obvious effect of lowering FAIMS resolution, this does not alter the type of peptides being identified, indicated by very similar precursor m/z range and distribution over the entire retention time (Error! Reference source not found. **E)**.

## Discussion

This study demonstrates that low-resolution FAIMS significantly increases peptide identifications from low-load samples. By broadening the compensation voltage window, low-resolution FAIMS enhances ion transmission, leading to a notable improvement in peptide identifications ranging from 250 pg to 16 ng HeLa digests. Although, to a lesser extent, we demonstrated that this adaptation is also beneficial for single-cell samples, where enhanced sensitivity is a crucial benefit from the here proposed strategy for detecting low-abundance proteins that might otherwise be missed and improve the quality of quantification crucial for all biological applications in the single cell proteomic field.

Moreover, we found that employing “low-resolution” FAIMS increases the ion counts of the measurements, both in MS1 and MS2 which has a positive effect on the quantitative precision. We consistently observed slightly lowered CoVs of samples measured with a lower outer electrode temperature.

The findings offer a practical optimization strategy for FAIMS-based low-input proteomics workflows. By simply adjusting the FAIMS resolution, researchers can achieve improved results without the need for additional hardware or changing data processing. This makes low-resolution FAIMS a cost-effective and straightforward enhancement for existing low-load proteomics setups.

## Methods

### Cultivation of H460 cells

Cells were cultured at 37 °C in a humidified atmosphere at 5% CO_2_. H460 cells were grown in RPMI 1640 medium. Cell medium was supplemented with 10% fetal bovine serum (FBS; 10270, Fisher Scientific), 1× penicillin–streptomycin (P0781-100ML, Sigma-Aldrich), 100× l-glutamine (200 mM, 250030-024, Thermo Scientific) and 1 mM sodium pyruvate (for RPMI only; 4275, Sigma-Aldrich). Cells were grown to around 75% confluency before trypsinization with 0.05% Trypsin-EDTA (25300-054, Thermo Scientific), followed by washing three times with PBS. Cells were resuspended in PBS at a density of 200 cells/μl for isolation using the cellenONE (Cellenion).

### Single cell proteomic sample preparation

Individual ell isolation, lysis and digestion were performed within a 384-well plate (Eppendorf twin.tec PCR Plate 384 LoBind) using the cellenONE robot based on our previously published One-Pot protocol.^16^ The cellenONE robot was operated using its control software (v2.0-1146). Cells were sorted into wells containing 1 µl of master mix (0.2% DDM (D4641-500MG, Sigma-Aldrich), 100 mM triethylammonium bicarbonate (TEAB; 17902-500ML, Fluka Analytical), 3 ng µl^–1^ trypsin (Trypsin Gold, V5280, Promega), 0.01% enhancer (ProteaseMAX, V2071, Promega) and 1% DMSO). Humidity and temperature were controlled at 50% and 15 °C during cell sorting. H460 cells were isolated at a given diameter of 20–35 µm. The maximum elongation was set to 1.6. Cell lysis and protein digestion were performed at 40 °C and 70% relative humidity for 30 min before an additional 500 nl of 3 ng μl^–1^ trypsin was added. After lysis and digestion, 3.0 μl of 0.1% TFA was added to the respective wells for quenching and storage at −20 °C. For LC–MS/MS analysis, samples were directly injected from the 384-well plate.

### Diluted bulk digests

HeLa (Thermo Scientific, Pierce HeLa Protein Digest Standard, 88328) peptides were dissolved in 0.1% TFA and 0.015% DDM (D4641-500MG, Sigma-Aldrich) at a concentration of 5 ng µl^–1^ for injection into the LC–MS system. Injection amounts were adopted from 0.05 µL to 3.2 µL to inject sample amounts ranging from a single cell equivalent (250 pg) - 16 ng respectively.

### LC–MS analysis

Samples were analyzed using the Thermo Scientific Vanquish Neo UHPLC system. Thermo Tune software version 1.1.477.46 or higher was used to acquire data. Peptides were loaded on a trapping column (Thermo Fisher Scientific, PepMap C18, 5 mm × 300 μm i.d., 5 μm particles, 100 Å pore size) using 0.1% TFA as the mobile phase. Peptides were separated on an Aurora Rapid^®^ 8×75 XT C18 nanoflow UHPLC column with an integrated emitter (Ion Optics) at 50 °C. Peptide separation was performed at 50–80 SPD with details provided in Supplementary Table 1 and with all single-cell measurements performed at 80 SPD.

For MS measuring, the Orbitrap Astral mass spectrometer (Thermo Scientific) equipped with a FAIMS Pro Duo interface (Thermo Scientific) and an EASY-Spray source was coupled to the LC. A compensation voltage of –48 V was used. An electrospray voltage of 1.9 kV was applied for ionization and adopted to slightly higher voltages for aged emitters to ensure spray stability.

MS method settings used were based on previous work.^17^ In brief, MS^1^ spectra were recorded using the Orbitrap analyzer at a resolution of 240,000 from *m*/*z* 400 to 900 using an automated gate control (AGC) target of 500% and a maximum injection time of 100 ms. Non-overlapping isolation windows of *m*/*z* of 20 were used for MS^2^ in DIA mode using the Astral analyzer. A scan range from an *m*/*z* of 400 to 800 was chosen. The precursor accumulation time was at a maximum of 40 ms and the AGC target was set to 500%.

### Data analysis

All raw data were analyzed using Spectronaut (version 20.1, Biognosys). Data was searched using directDIA+ in method evaluation mode allowing for cross normalization within each tested condition. Quantification was performed at the MS^1^ level. Unless otherwise indicated, default settings were used, with carbamidomethylation of cysteines as a static modification was removed for single cell searches as no alkylation step was performed. Searches were performed against the human proteome (UniProt proteome UP000005640, reviewed, 20,408 protein entries, downloaded 4^th^ of August 2023) and the CRAPome^18^ (118 protein entries). If not otherwise stated, FDR filtering was based on default settings of Spectronaut and at 1% at the protein level. Post processing was done using R and Graph Pad Prism (v8.1.1). For PCA no additional imputation or normalization was applied.

The LC peak total ion counts from 250pg HeLa 50SPD runs were extracted using Skyline (v25.1.0.142)^19^ using the ion count report as used by Hsu et al.^20^

## Acknowledgments

This work was supported by the infrastructure funding 4^th^ call 2022/01 (AT-SCP) of the Austrian Research Promotion Agency (FFG) The authors thank the group of Josef Penninger at IMBA for providing H460 cells. For the purpose of open access, the authors have applied a CC BY public copyright license to this submission.

## Author Contributions Statement

DGH and MM conceptualized the study, designed experiments, performed data analysis, and wrote the manuscript. MM performed all experiments. KM supervised the study. All authors revised and agreed on the manuscript.

## Supplemental Tables and Figures

**Supplemental Figure 11:**
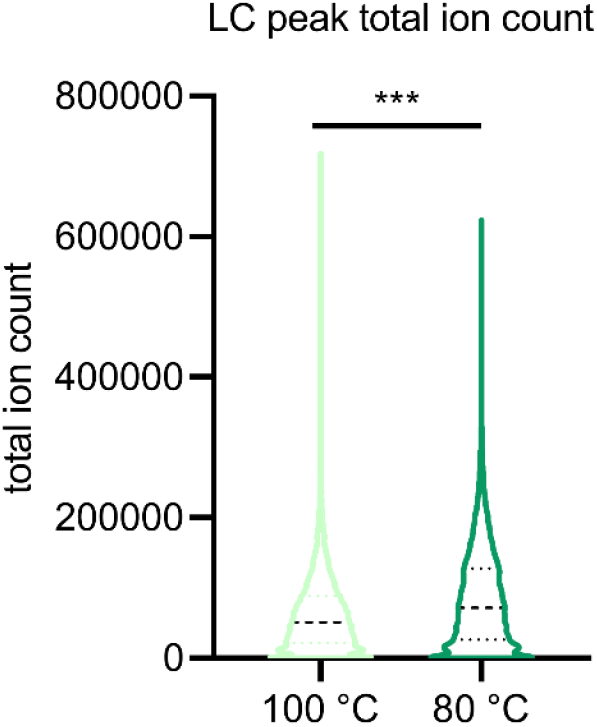
LC peak total ion count. Box plot of the LC peak total fragment ion counts of a 250pg HeLa 50SPD run with standard(left) and low-resolution (right) FAIMS setting. For statistical comparison an unpaired two tailed students t test was performed, with *** for p<0.001,

**Supplemental Figure 22:**
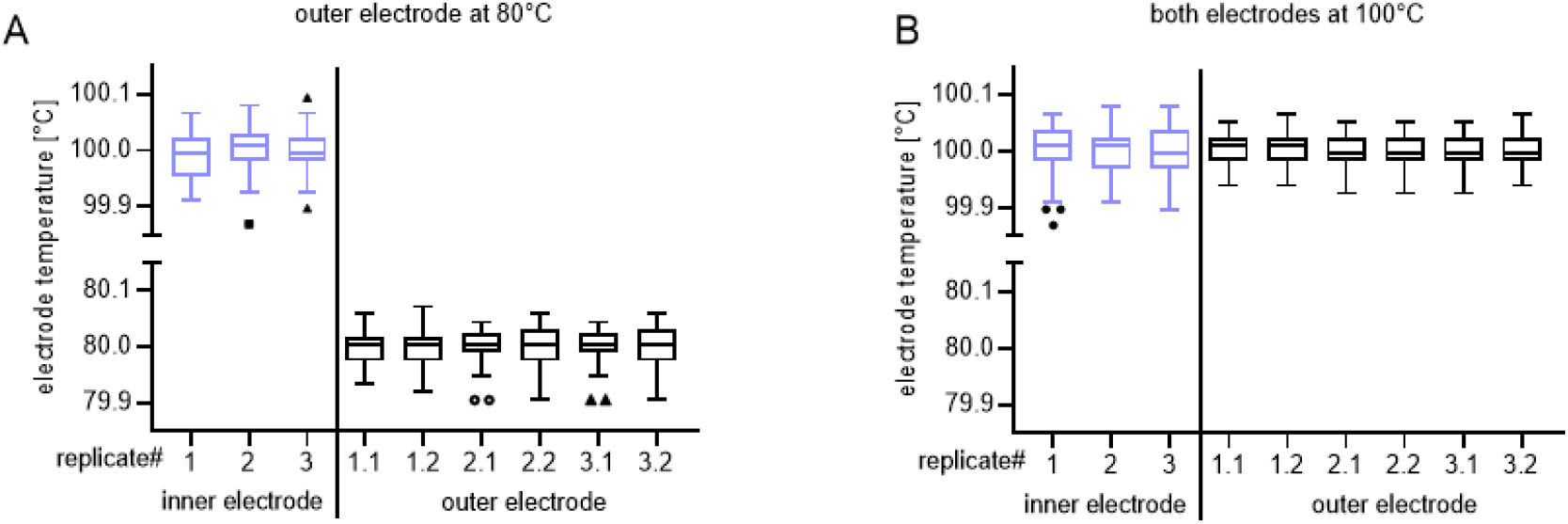
Temperature stability of inner and outer electrode. (**A**) Low-resolution mode with the outer electrode temperature set to 80°C. (**B**) Standard resolution mode. Each boxplot shows the temperature distribution of each electrode during an entire 80SPD run. Temperature of the outer electrode is measured at two points in the FAIMS electrode assembly, yielding double the datapoints compared to the inner electrode temperature measurements (specified as x.1 and x.2).

**Supplemental Table 1:** LC gradient details.

|  | 50 SPD |  | 80 SPD |  |
| --- | --- | --- | --- | --- |
| TIME [MIN] | flow [nl/min] | % buffer B | flow [nl/min] | % buffer B |
| **0** | 450 | 1 | 1000 | 6 |
| **0.1** | 450 | 4 | - | - |
| **0.5** | - | - | 1000 | 10 |
| **1.0** | - | - | 250 | 12 |
| **1.9** | 450 | 12 | - | - |
| **2** | 200 | - | - | - |
| **10.5** | - | - | 250 | 37 |
| **12** | 200 | 22.5 | - | - |
| **12.5** | - | - | 250 | 56 |
| **13** | - | - | 1000 | 99 |
| **14** | - | - | 1000 | 99 |
| **19.5** | 200 | 40 | - | - |
| **20** | - | - | - | - |
| **22** | 300 | 99 | - | - |
| **25** | 300 | 99 | - | - |

