## Supplementary figures and images for "Low-resolution FAIMS for increased peptide coverage in low-load and single-cell proteomics"

### Supplemental Figure 1

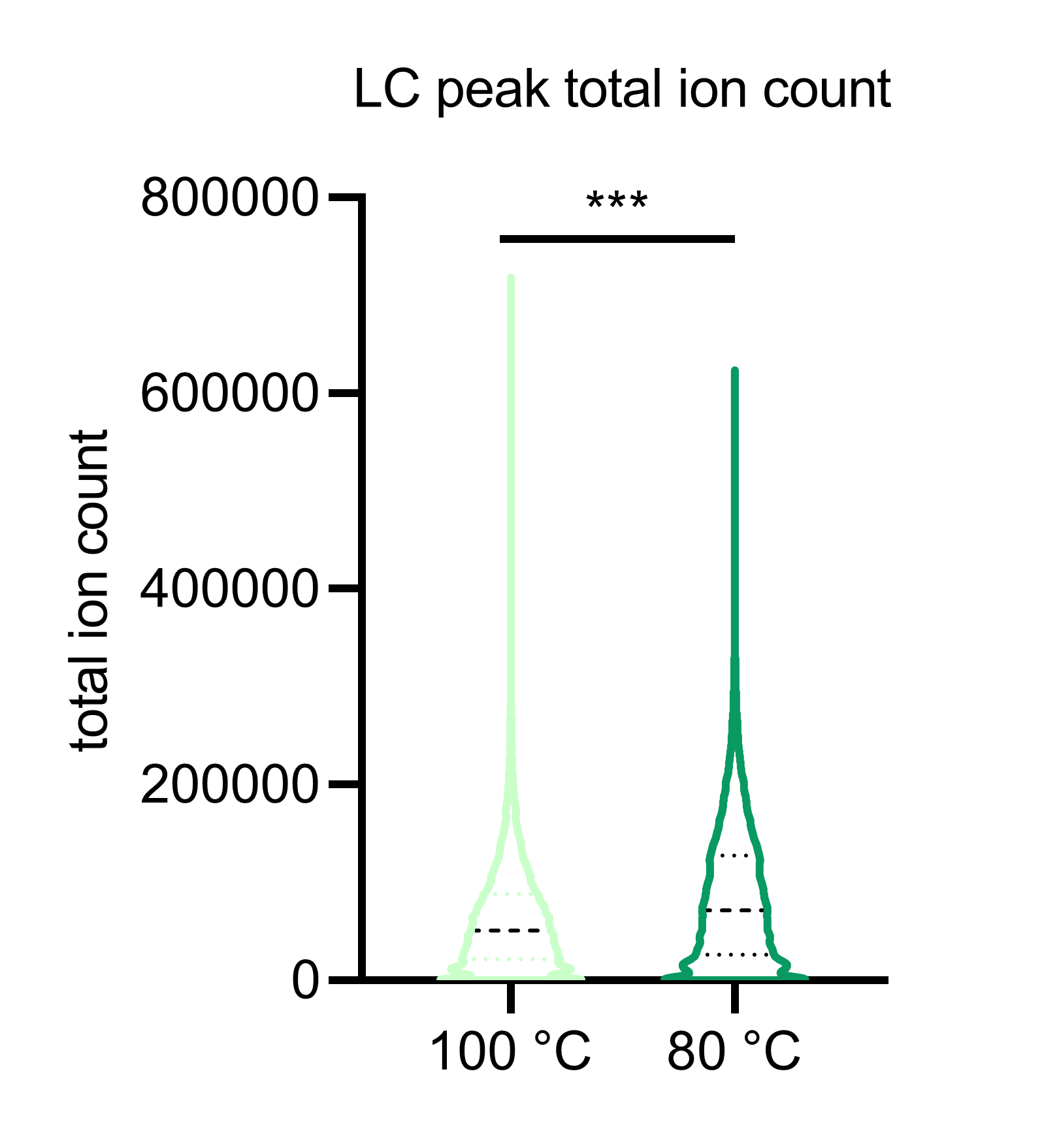

### Supplemental Figure 2

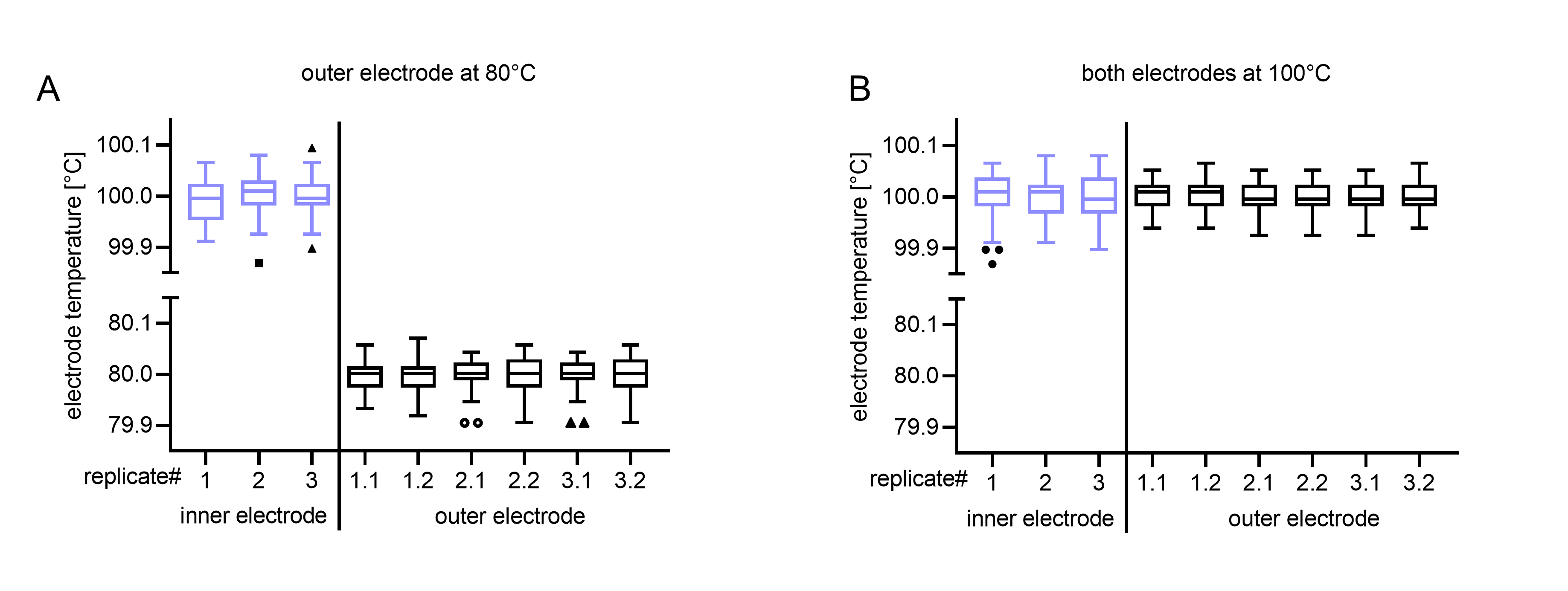
